# Putrescine acts as a signaling metabolite in the transition from nodulation to nitrogen fixation in *Rhizobium phaseoli*

**DOI:** 10.1101/2024.01.29.577838

**Authors:** Ericka M. Hernandez-Benitez, Esperanza Martínez-Romero, José Luis Aguirre-Noyola, Daniela Ledezma-Tejeida

**Author notes:** To whom correspondence must be addressed.

## Abstract

Growth of the common bean plant *Phaseolus vulgaris* is tightly linked to its symbiotic relationship with diverse rhizobial species, particularly *Rhizobium phaseoli*, an alphaproteobacteria that forms root nodules and provides high levels of nitrogen to the plant. Molecular cross-talk is known to happen via plant-derived metabolites, but only flavonoids have been identified as signals. Flavonoids are transported inside the bacteria, where they signal the NodD regulator to elicit nodulation. Although seven other regulators are known to be involved, our knowledge of the regulatory mechanisms underlying the nodulation, and nitrogen fixation processes is limited, and the signals recognized by regulators are mostly unknown. Here, we identified 75 transcription factors in *R. phaseoli* genome through sequence conservation from *Escherichia coli*, and assembled a transcriptional regulatory network comprising 24 regulators, and 652 target genes. We identified the interactions relevant to nodulation via gene expression, and signaled out putrescine as a signaling metabolite. We propose a model where putrescine acts as a switch on the transition from nodulation to nitrogen fixation via the dual transcription factor PuuR, and its regulation of the *nodI* and *nifU2* genes.

**Importance:** This study provides new insights into the molecular cross-talk between *Phaseolus vulgaris* and *Rhizobium phaseoli*, which is crucial for the development of alternative cropping strategies, including biopesticides and biofertilizers. In addition, we present the first transcriptional regulatory network systematically assembled for *Rhizobium phaseoli*, which opens new avenues of research in the relationship between the transcriptional regulation and metabolism of these organisms.

## BACKGROUND

*Rhizobium phaseoli* plays a significant role in leguminous plants’ growth, development, and health because of its ability to establish a symbiotic relationship through the root nodules (1–3). Within this specialized structure, the host plants provide C_4_-dicarboxylate and phosphate to the symbionts, which use them to fix atmospheric nitrogen into ammonium, a nitrogen source for the host plants (4). The symbiont life cycle comprises two stages: nodulation and nitrogen fixation. In the first, nodulation genes, commonly known as *nod* genes, are expressed to initiate the development of root nodules. In the nitrogen fixation stage, *nif* and *fix* genes are expressed to control the synthesis of nitrogenase complex proteins and to provide respiration and electron supply to nitrogenase, respectively (5).

In the past three decades, the utilization of plant-associated symbionts as biofertilizers and biopesticides has been proposed as the best alternative for low-cost, eco-friendly, and sustainable protection of crops, compared to agrochemicals (6). Nevertheless, to successfully develop biopesticides based on beneficial microorganisms, it is crucial to understand the symbiotic interaction, including the regulatory mechanisms and signaling molecules involved in plant-symbiont signaling (6).

One of the best-known regulatory mechanisms of the nodulation process is the activation of the *nodABC* operon through flavonoid signaling (7). After root exudation of these secondary metabolites by legumes, they are imported into *Rhizobium* bacteria, where they interact with the Transcription factor (TF) NodD. In turn, NodD promotes the activation of the *nodABC* operon, and the resulting synthesis of Nod Factors (NFs) (7, 8), lipooligosaccharides whose available functional groups have a role in host-specificity. NFs are exported by the bacteria to signal the plant and initiate the nodulation process (9). Other TFs involved in the nodulation process are NodV, NodW, and NolA, but they are not present in all *Rhizobium* species (8), and their signaling metabolites are unknown. Regulators of nitrogen fixation are the two-component FixL-FixJ, NifA, and the FNR/CRP family of TFs that include FixK and FnrN (10, 11). Here, low oxygen activates the FixL kinase, which in turn activates the FixJ regulator, resulting in FixK and FnrN activation of *fix* genes, and NifA-mediate activation of *nif* genes (11).

Similarities have been reported between *fnr*-like genes present in *Rhizobiaceae*, and the FNR TF in *E. coli* that regulates genes involved in anaerobic respiration (10–12). FixK from *Bradyrhizobium japonicum*, a known paralogue of FNR shares 28% of sequence identity (13), while FnrN another member of the FNR/CRP family in *Rhizobium leguminosarum*, shares an identity of 26% to FNR (12). Both TFs have been shown to also conserve their DNA-binding capabilities along with the DNA sequence they recognize, called the TF-binding site (TFBS), and both can complement an *E. coli* strain lacking FNR by regulating its target genes (TGs) (12, 14). Likewise, *E. coli* FNR can complement an *R. leguminosarum* mutant lacking FnrN by activating the *fixNOQP* operon in an oxygen-dependent manner (12). Along with the sequence conservation of a Cys-cluster that can detect fluctuations in oxygen level, the complementation of FnrN by *E. coli* FNR supports the evolutionary preservation of the signal stimulus (12).

Given this high level of conservation, we sought to expand our limited knowledge of signals mediating the symbiosis process by identifying other conserved TFs in *Rhizobium phaseoli*. We identified 75 *E. coli* TFs with sequence conservation in *R. phaseoli*, of which 58 had a reported DNA-binding domain, and 84% had it conserved. We assembled a network of TF-gene interactions by scanning the upstream regions of *R. phaseoli* promoters, resulting in 787 interactions involving 24 TFs and 652 TGs. Finally, we combined three levels of independent evidence to identify TFs regulating genes involved in the symbiosis process. We propose PuuR as a TF mediating the nodulation and nitrogen fixation processes and its binding metabolite putrescine as a signal in the transition from nodulation to nitrogen fixation.

## RESULTS

### Seventy-five *E. coli* Transcription Factors are conserved in *R. phaseoli*

To identify orthologous TFs (oTFs) in *R. phaseoli*, we used the well-characterized *E. coli* Transcriptional Regulatory Network (TRN), which includes 304 TFs – 108 predicted, and 196 experimentally characterized (15) – and 1823 TGs (16). Even though *E. coli* does not participate in nodulation or nitrogen fixation processes, we expect the DNA-binding mediated regulatory function of the TFs to be conserved, albeit not necessarily the function of the TGs because non-coding sequences (TF-binding sites) diverge faster than coding sequences (DNA-binding motifs) (17). We looked for Bidirectional Best Hits (BBHs) using protein sequences (BLASTp). We considered a pair of TFs orthologous if they had an identity of ≥ 30% and an e-value ≤ 1x10^−3^ based on literature (18). Although “orthologue” had its original definition as an evolutionary relationship based on speciation, that is not our claim. Here, we will use the more common definition of genes with conserved functions, for which sequence similarity has been shown to be the most effective method (19). From the 4285 *E. coli* coding genes, we identified 1266 orthologous genes in *R. phaseoli*, of which only 75 were oTFs (**Figure 1, Supplementary Figure 1**). On average, the oTFs have a length of 277 amino acids in *E. coli* and 275 in *R. phaseoli* (**Supplementary Figure 2**). Nine TFs (AlaS, DecR, IhfA, PgrR, PutA, RtcR, RutR, SoxR, and YoeB) have an identity and query coverage ≥ 50% (**Figure 1**, upper right quadrant of both dot plots), with PgrR being the TF with the highest query coverage (99.33% and 100%) and SoxR the TF with the highest identity (67.83%).

**Figure 1.**
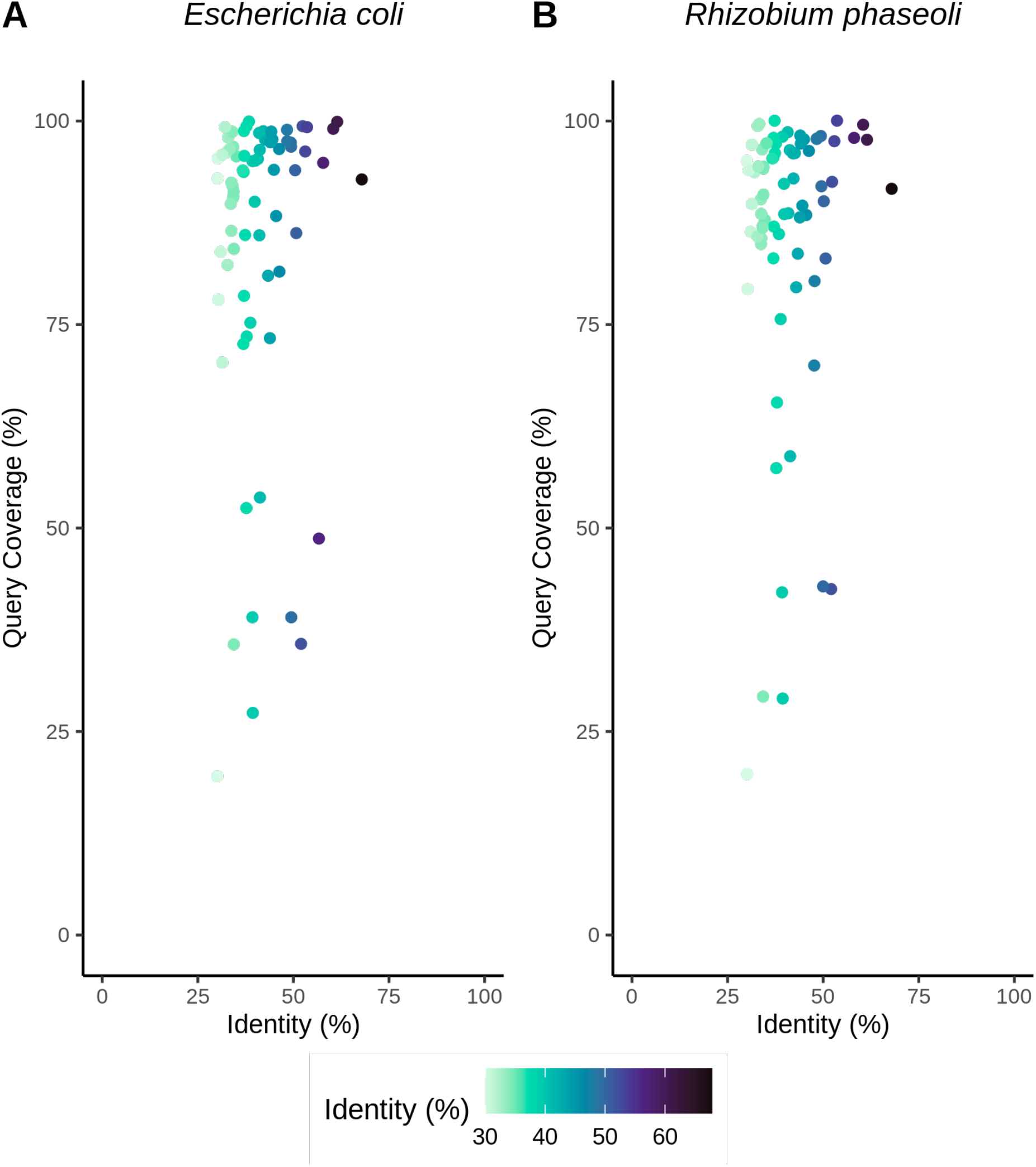
Identity and coverage of orthologous TFs. Percentages are shown using the (A) *E. coli* or (B) *R. phaseoli* genes as reference. Each point represents a TF. The highest-scoring TFs were AlaS, DecR, IhfA, PgrR, PutA, RtcR, RutR, SoxR, and YoeB.

### DNA-binding motifs are highly conserved in orthologous TFs

To support the conservation of TFBS-recognition and DNA-binding capacities on oTFs, we assessed the conservation of their reported protein motifs. Motif annotation for all oTFs was retrieved from the Ecocyc database given its high level of curation (20) (**Supplementary Table 1**). Annotation includes 144 motifs for 62 TFs. On average, each TF had 2.32 motifs reported. Thirteen TFs had no motifs reported and were excluded from further analyses. For the remaining 62 TFs, a motif was considered conserved if it had an identity ≥ 30% in the alignment with the oTF in *R. phaseoli*. We classified the oTFs into three categories, depending on their level of motif conservation: (1) ‘All conserved motifs’ when every reported motif in the protein sequence was conserved. (2) ‘At least one conserved motif’ when one or more of the reported motifs, but not all, were conserved. (3) ‘No conserved motifs’ when none of the reported motifs bypassed our threshold for conservation. Additionally, we examined the types of motifs found in each category.

**Table 1.** Summary of the TFs in the *R. phaseoli* Transcriptional Regulatory Network. Each row corresponds to a TF in the Transcriptional Regulatory Network. TF name: TF canonical name in *E. coli*; TF locus-tag: Unique gene identifier in *E. coli;* DNA-binding region/Conserved region: indicate whether the reported motif in Ecocyc is conserved or not. If the motif is conserved, the motif coordinates are displayed, otherwise, an NA is shown; Effector name: The metabolite(s) known to bind in *E. coli*; Number of TGs: Number of Target Genes per TF in the Transcriptional Regulatory Network.

| TF name | TF locus-tag | DNA-binding region | Conserved region | Effector name | Number of TGs |
| --- | --- | --- | --- | --- | --- |
| AcrR | b0464 | 33-52 | 10-70 | Proflavin, ethidium | 29 |
| ArgP | b2916 | 21-40 | NA | L-lysine, L-arginine | 41 |
| BaeR | b2079 | 131-234 | 12-125 | NA | 24 |
| CytR | b3934 | 12-31 | 10-64 | cytidine | 32 |
| Dan | b3060 | 23-42 | NA | L-tartrate | 14 |
| ExuR | b3094 | 35-54 | 7-75 | D-glucuronate, D-galacturonate | 20 |
| FNR | b1334 | 197-216 | 164-237 | [2Fe-2S] oxidized, [4Fe-4S] reduced | 36 |
| GalR | b2837 | 4-23 | NA | D-galactose | 18 |
| GcvA | b2808 | 23-42 | 6-63 | glycine, Purine | 24 |
| GntR | b3438 | 8-27 | 6-60 | D-gluconate | 46 |
| IscR | b2531 | 28-51 | NA | a [2Fe-2S] iron-sulfur cluster | 39 |
| Lrp | b0889 | 31-50 | 12-73 | L-leucine | 35 |
| Nac | b1988 | 18-37 | 1-58 | NA | 35 |
| NarP | b2193 | 171-190 | NA, 8-124 | NA | 58 |
| NsrR | b4178 | 28-51 | 2-129 | nitric oxide, [2Fe-2S] reduced, [2Fe-2S] oxidized | 28 |
| NtrC | b3868 | 445-464 | NA, 140-369 | NA | 17 |
| PurR | b1658 | 4-23, 48-56 | 2-56 | Guanine, hypoxanthine | 20 |
| PuuR | b1299 | 23-42 | 12-66 | putrescine | 60 |
| RhaS | b3905 | 191-212, 239-262 | NA | L-rhamnose | 43 |
| Rob | b4396 | 73-96, 25-46 | NA | NA | 15 |
| RutR | b1013 | 39-58 | NA | uracil, thymine | 28 |
| SlyA | b1642 | 49-72 | NA | NA | 43 |
| SoxR | b4063 | 14-33 | 11-79 | [2Fe-2S] reduced, [2Fe-2S] oxydized | 43 |
| UlaR | b4191 | 20-39 | 3-58 | L-ascorbate 6-phosphate | 39 |

The most common category was oTFs with all their motifs conserved, which included 80.65% (50/62) oTFs, 11.29% (7/62) fell under the ‘At least one conserved motif’ categories, and only 8.06% (5/62) had no conserved motifs, (**Supplementary Figure 3A**). “DNA-binding regions” and “Conserved regions” were the main types of conserved motifs in the ‘All conserved motifs’ category, with 45.95% (51/111) and 47.75% (53/111), respectively (**Supplementary Figure 3B**), followed by 5.41% (6/111) for Nucleotide-Phosphate-Binding-Region and 0.9% (1/111) for Zn-Finger-Region.

Because DNA-binding motifs are characteristic of functional TFs, we further analyzed their conservation within oTFs. Of the 75 oTFs, 58 have at least one reported “DNA-binding region” motif, and 49 (84%) had their “DNA-binding region” motif conserved (**Figure 2**). The conserved “DNA-binding region” motifs have, on average, an identity of 49.89% and a coverage of 98.78%. The high percentage of conserved motifs supports the conservation of protein function and the role of oTFs as transcriptional regulators in *R. phaseoli*.

**Figure 2.**
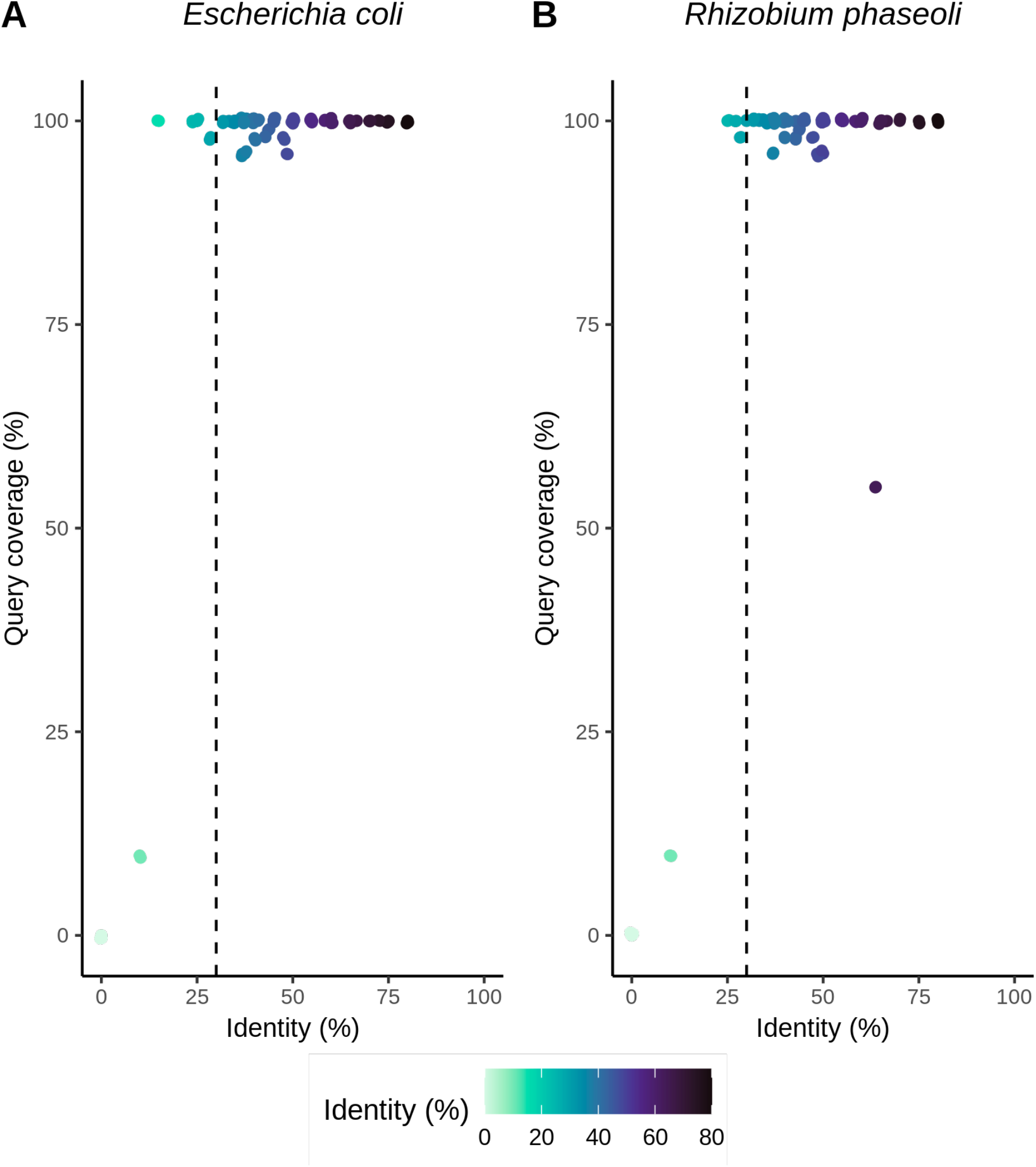
Identity and coverage of DNA-binding regions. Percentages are shown with (A) *E. coli* or (B) *R. phaseoli* genes as reference. Each point represents a DNA-binding region of a TF. Identity cut-off values for the motif to be considered conserved are shown in dashed lines.

### Inference of *R. phaseoli* transcriptional regulatory network using orthologous TFs

The next goal was to assemble a TRN for *R. phaseoli* from the identified oTFs. The identification of regulated genes is commonly done by scanning the genome for known TFBSs. Given the high conservation of DNA-binding motifs between *E. coli* and *R. phaseoli*, we used the available Position-Specific Scoring Matrices (PSSMs) from *E. coli* to scan the *R. phaseoli* genome under the assumption that non-coding sequences diverge faster than coding regions, and therefore an oTF is likely to recognize a very similar TFBS, but the site will not necessarily be upstream of an orthologous TG. To increase the reliability of the TRN, we extracted the subset of oTFs with a sequence coverage of 100% and an identity ≥30% in their “DNA-Binding Region” motif. Forty-one oTFs passed the more stringent cutoff, and 25 had an available PSSM in the RegulonDB database. *R. phaseoli* upstream gene sequences were defined from position -400 to +50 relative to the Transcription Start Site (TSS) and were scanned using the matrix-scan tool (21) from the Regulatory Sequence Analysis Tools (RSAT) suite (**Figure 3**). We selected a p-value cut-off ≤ 1e-5, which indicates that we expect one match by chance for each of the 25 PSSMs every 100 kb. Since the scanned sequences are 450 bp in length, the probability of having a match by chance is 4.5e-3 for each PSSM. Even considering a multiple-test correction for the 25 PSSM, the FDR is 0.1125, lower than 1 expected false positive. We assumed an interaction when a match between a PSSM and an upstream sequence was present.

**Figure 3.**
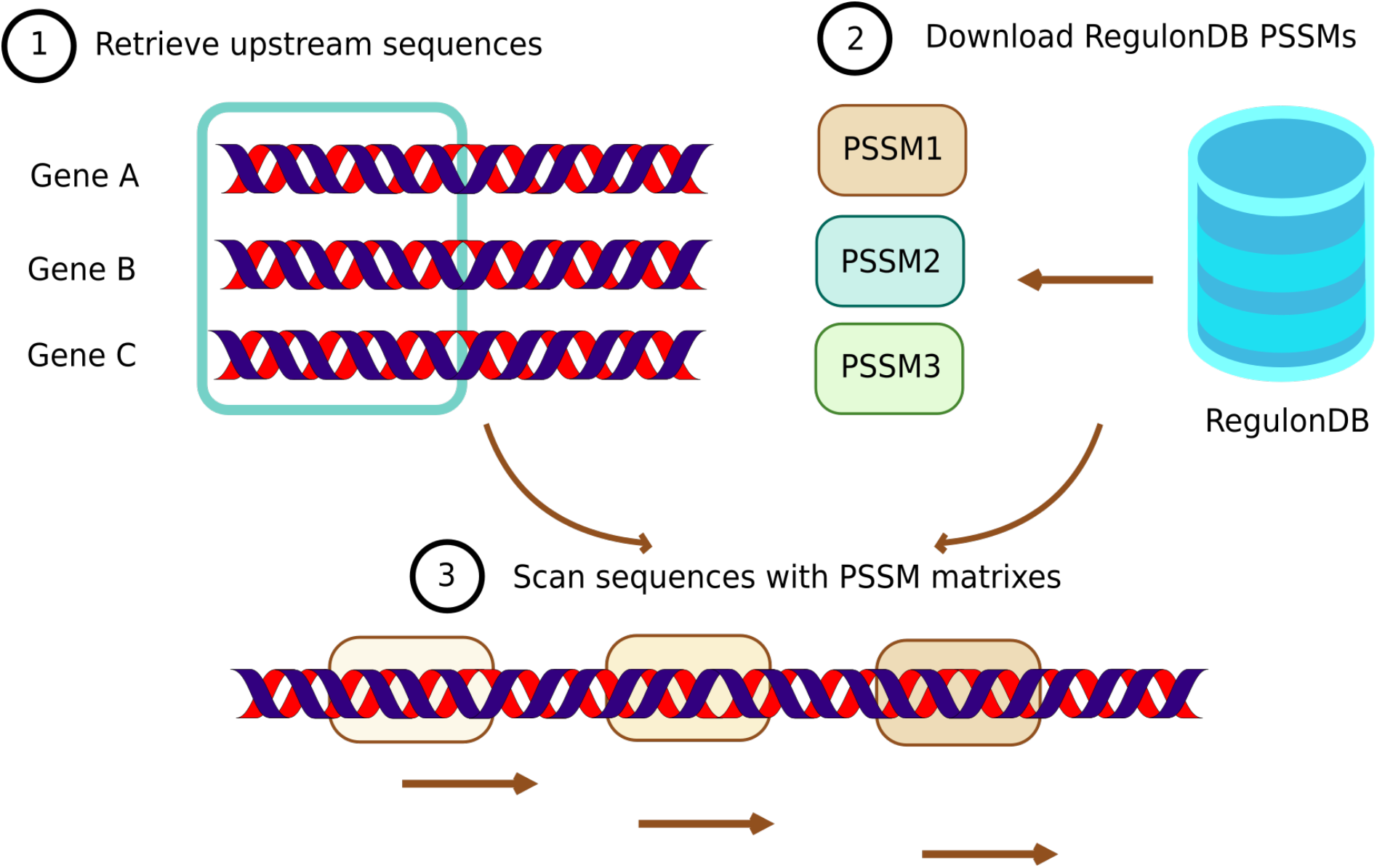
Construction of the *R. phaseoli* Transcriptional Regulatory Network. Upstream sequences from all *R. phaseoli* genes were scanned with orthologous TFs’ Position-Specific Scoring Matrices, obtained from RegulonDB, using the scan-matrix software (21).

The resulting TRN consisted of 787 TF-TG interactions between 24 TFs and 652 TGs, with each TF regulating 32 TGs on average (**Figure 4, Table 1**). Only 110 (16.87%) TGs have an orthologue in *E. coli*, which was consistent with our expectation of TFs recognizing a similar TFBS, but regulating different genes. Furthermore, only 5 TF-TG interactions are conserved in both *E. coli* and *R. phaseoli*, of which three are TFs (FNR, PurR, and RutR) regulating their own coding-gene, a self-regulation dynamic that has been shown to be positively selected in genetic circuits (22).

**Figure 4.**
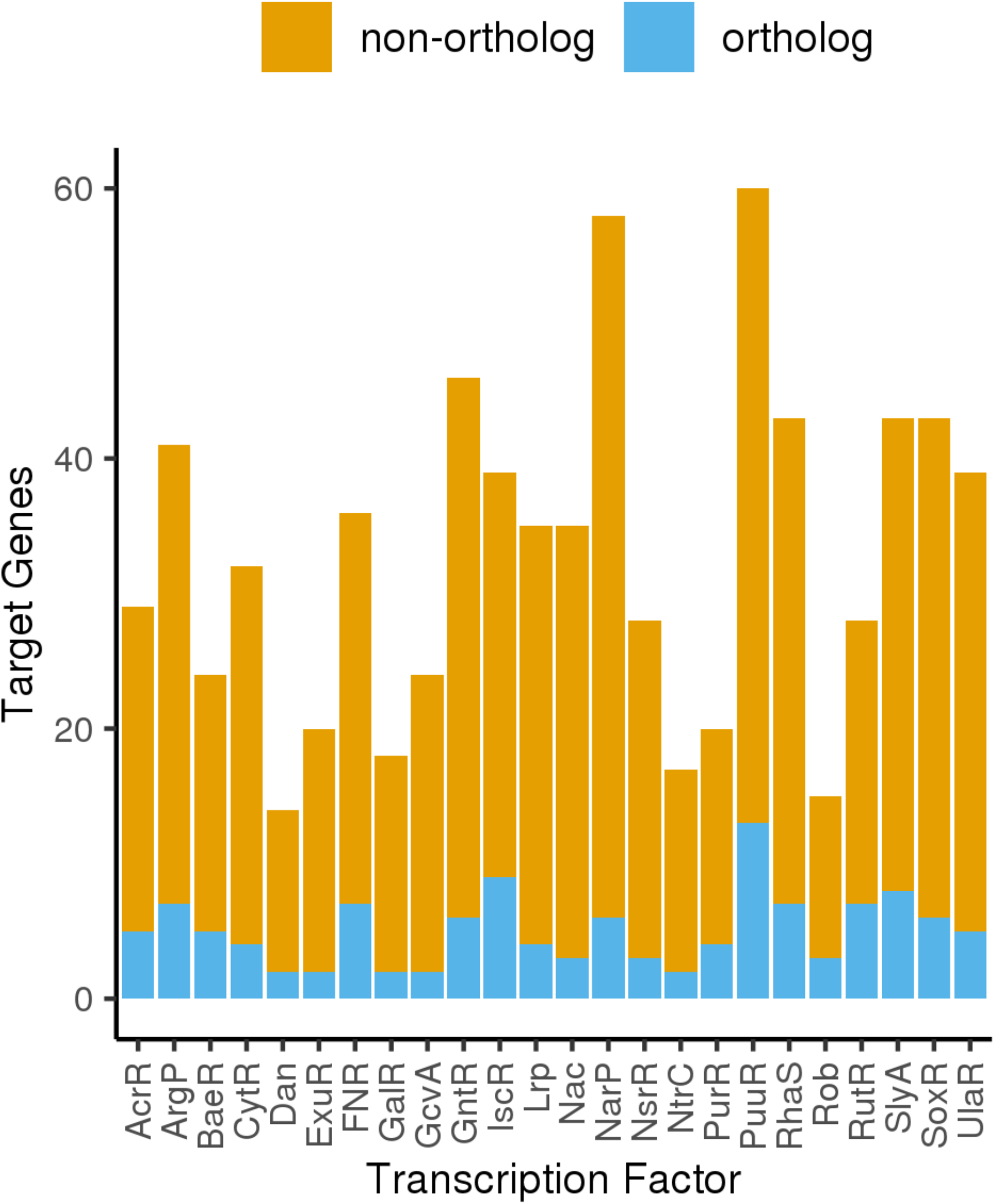
*R. phaseoli* Transcriptional Regulatory Network. Number of TGs (y-axis) per TF (x-axis). Each bar shows the proportion of orthologous and non-orthologous TGs.

We identified an orthologous relationship between the *E. coli* TF FNR and the *R. phaseoli* TF FixK, which had been reported to be conserved in other nitrogen-fixing bacteria belonging to the family *Rhizobiaceae* including *Sinorhizobium meliloti* and *Rhizobium leguminosarum* (10). This relationship showed an identity of 30.35% and a query coverage of 78% and 79.42% in the BBH. Moreover, we identified a previously reported regulatory interaction in *Sinorhizobium meliloti, Rhizobium leguminosarum*, and *Rhizobium etli* (10) between the TF fixK and the nitrogen fixation gene, *fixN*, in the reconstructed *R. phaseoli* TRN. To the best of our knowledge, efforts to identify TF-TG interactions in *Rhizobium* have been limited to *Rhizobium etli* and *Rhizobium leguminosarum* (23–25), so this TRN represents the first effort to systematically identify TF-TG interactions in *Rhizobium phaseoli*.

### PuuR is a regulator of nitrogen fixation in *R. phaseoli*

The high conservation of oTFs coupled with the low conservation of TF-TG interactions led us to hypothesize that some of the oTFs might regulate genes involved in Rhizobia-legume symbiosis. To identify these genes, we used two independent approaches. In the first approach, we searched in the literature for genes with experimental evidence (5, 8) of being involved in symbiosis, nodulation, or nitrogen fixation in *R. phaseoli* (**Supplementary Table 2**) under the rationale that oTFs regulating these genes would consequently have a role in the symbiotic process. As a second approach, we used a previously generated RNA-seq dataset of *R. phaseoli* grown in the presence of root exudates from Maize (*Zea mays*), common bean (*Phaseolus vulgaris*), and milpa system (i.e. maize-bean intercropping) under hydroponic conditions (26). Genes showing differential expression when compared to a control without root exudates were deemed as relevant for the symbiotic process. The overlap of the three layers of evidence – presence in the TRN, known function, and differential expression – yielded a gradient of confidence in the involvement of a gene in symbiosis.

We identified nine genes that show both evidence of function and evidence of gene expression: *nodA, nodB, nodC, nodS, nodT, nodZ, nifR, nifSch, nolO* (**Figure 5, Supplementary Figure 4**), and 11 genes with evidence of function and present in our reconstructed TRN: *nodD3, nifA, fnrN, fixA, fixG, fixGa, fixKa, fixN, fixNa, fnrNch*, and *nifU2*. The latter are regulated by 9 TFs: NarP, ArgP, CytR, GcvA, UlaR, ExuR, FNR, GntR, and PuuR, which are likely to regulate the symbiotic process (**Figure 5, Supplementary Table 3**). One gene, *nodI*, was present in the intersection of the three types of evidence and therefore was the most reliable result (**Figure 5**). In the TRN, *nodI*, is regulated by the oTF PuuR, which binds putrescine in *E. coli* (27).

**Figure 5.**
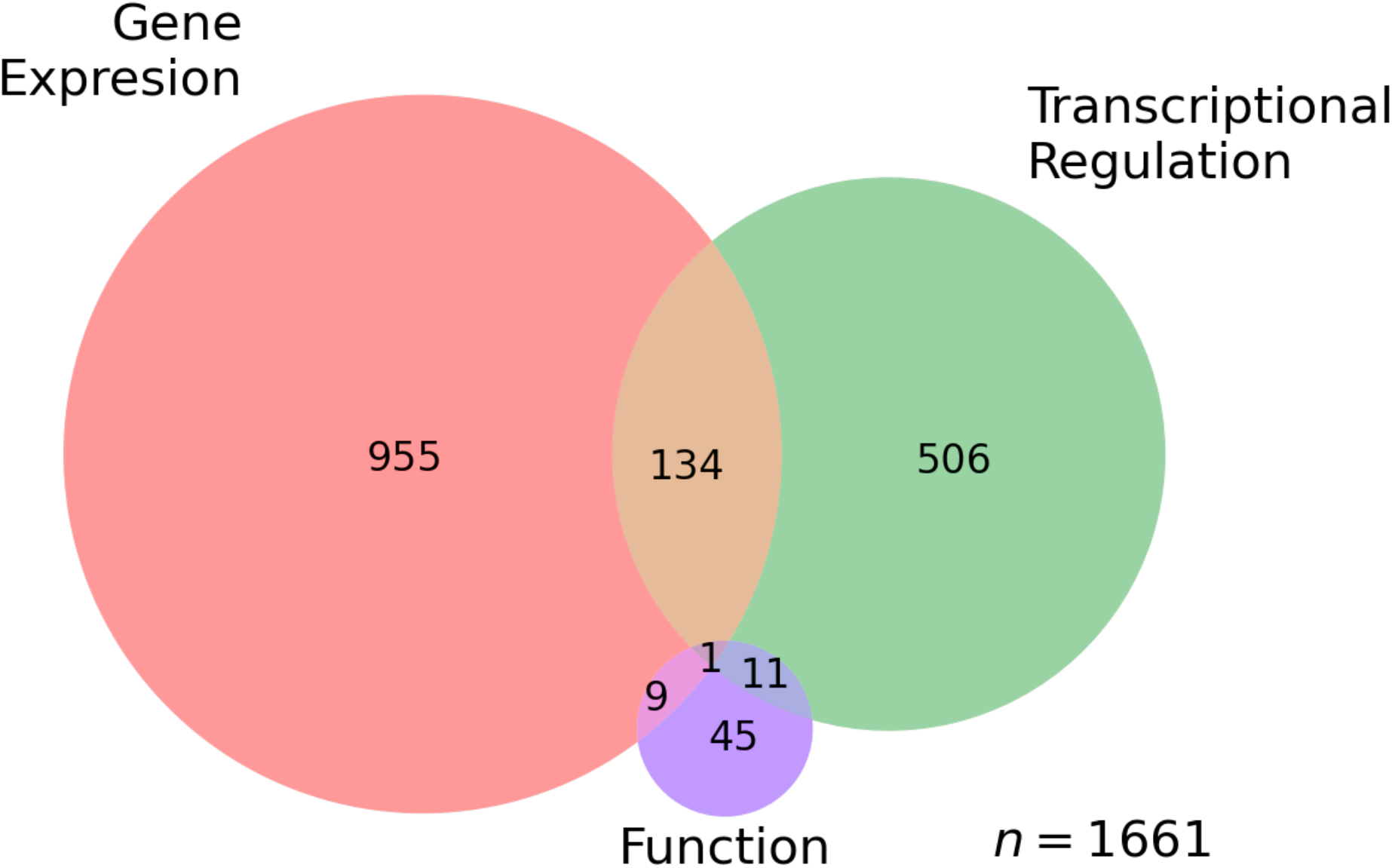
Levels of evidence. Levels of evidence supporting gene involvement in a symbiosis. Nine genes had evidence of function and gene expression; eleven genes had evidence of function and transcriptional regulation; and one gene had evidence of function, gene expression, and transcriptional regulation.

To add evidence of the PuuR-*nodI* interaction, we analyzed the similarity of the three-dimensional structure of the DNA-binding region motif of PuuR and its orthologue in *R. phaseoli*. Since no experimentally derived structures have been elucidated yet for either of these proteins (28), we obtained their predicted structure from the Alpha fold database (29). The DNA-binding region motifs of both TFs were aligned using the pairwise alignment tool from the RCSB PDB. An RMSD of 0.39 and a TM-score of 0.49 were obtained when aligning the DNA-binding region motifs, a significantly larger score than 0.2, the threshold indicating that proteins are unrelated. These results suggest that the motif conserves the same folding pattern and supports the conservation of TFBS recognition. We propose the PuuR-*nodI* interaction as a relevant regulatory interaction in the symbiotic process of *R. phaseoli*.

### PuuR is conserved in other nitrogen-fixing bacteria

To further support the functional relevance of PuuR in symbiosis, we evaluated its conservation among rhizobia and other closely-related bacteria. We identified orthologous sequences of *R. phaseoli* PuuR in seven *alpha*-proteobacteria from the *Hyphomicrobiales* order using BBH (**Supplementary Table 4**). We considered an oTF of PuuR if it had an identity percent ≥ 30% (**Supplementary Figure 5**). All species analyzed showed a conserved PuuR oTF with a length of 182 amino acids, a query coverage of 100%, and an identity ≥ 75% (**Figure 6**), indicating that the protein sequence is highly conserved. Furthermore, the annotation of the PuuR oTFs (30, 31) (**Supplementary Table 4**) is in agreement with our results, since they are annotated as “regulator” or “putrescine utilization regulator”. Both annotations are derived from the NCBI Prokaryotic Genome Annotation Pipeline (**PGAP**), which combines Hidden Markov Models-based gene prediction algorithms with protein sequence similarity search method as well as information of evolutionarily conserved proteins from Clusters of Orthologous Groups (COGs) and NCBI Prokaryotic Clusters to annotate genes (32). A maximum parsimony phylogenetic reconstruction of the PuuR oTFs with *E. coli* as an outgroup (**Figure 7** and **Supplementary Figure 6**) showed a clear division among the *Nitrobacteraceae* and *Rhizobiaceae* families. Subsequently, species are grouped by genus: *Sinorhizobium*, and *Rhizobium*. The PuuR phylogeny closely resembles a species phylogeny. All in all, our results support the activity of PuuR as a TF involved in nitrogen fixation.

**Figure 6.**
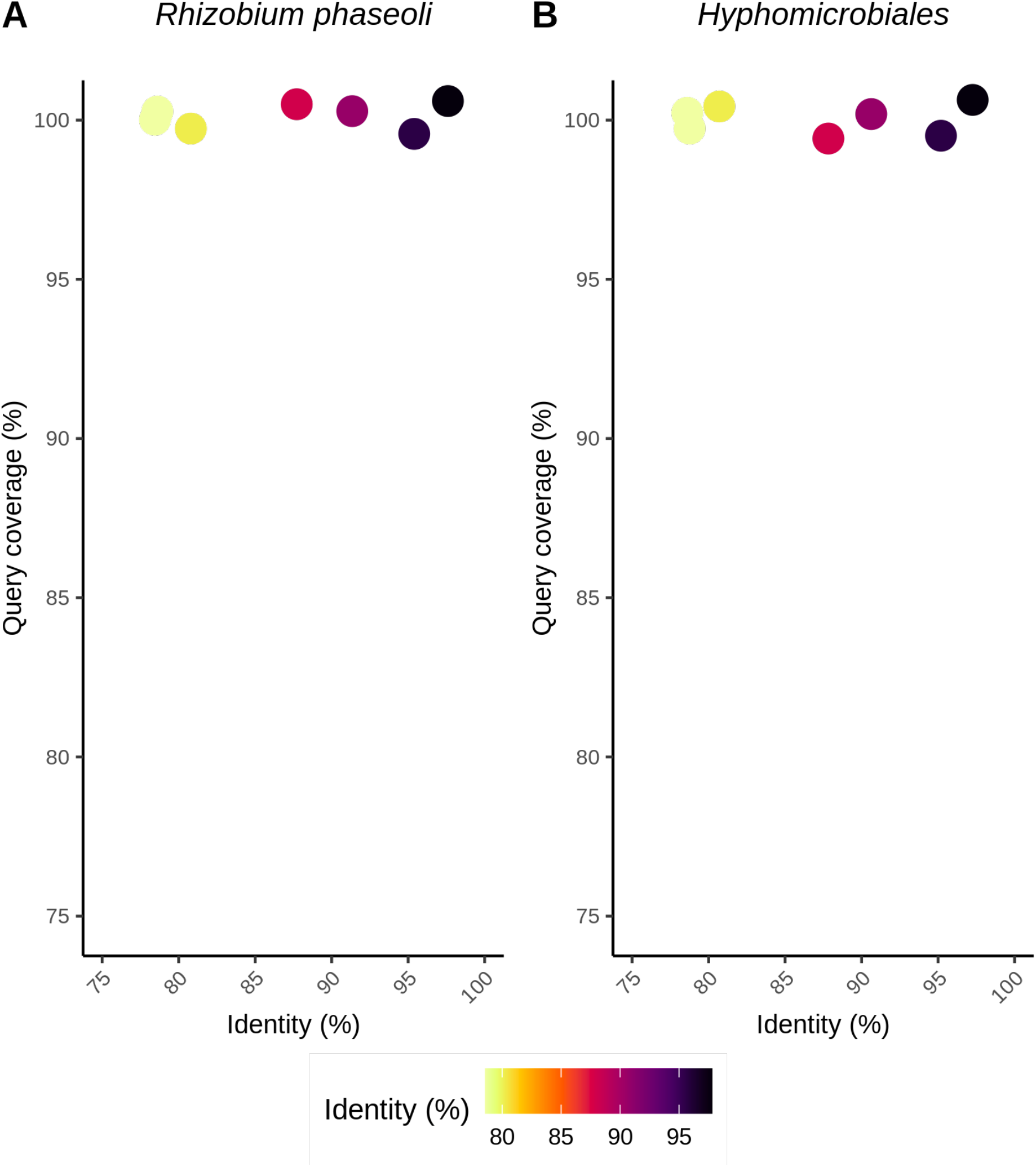
Identity and coverage of PuuR orthologous TFs. Orthologous PuuR regulatory proteins with (A) *R. phaseoli* or (B) *Hyphomicrobiales* proteins as reference. Each point represents an orthologous TF. No proteins were located below the 75% values in both plots.

**Figure 7.**
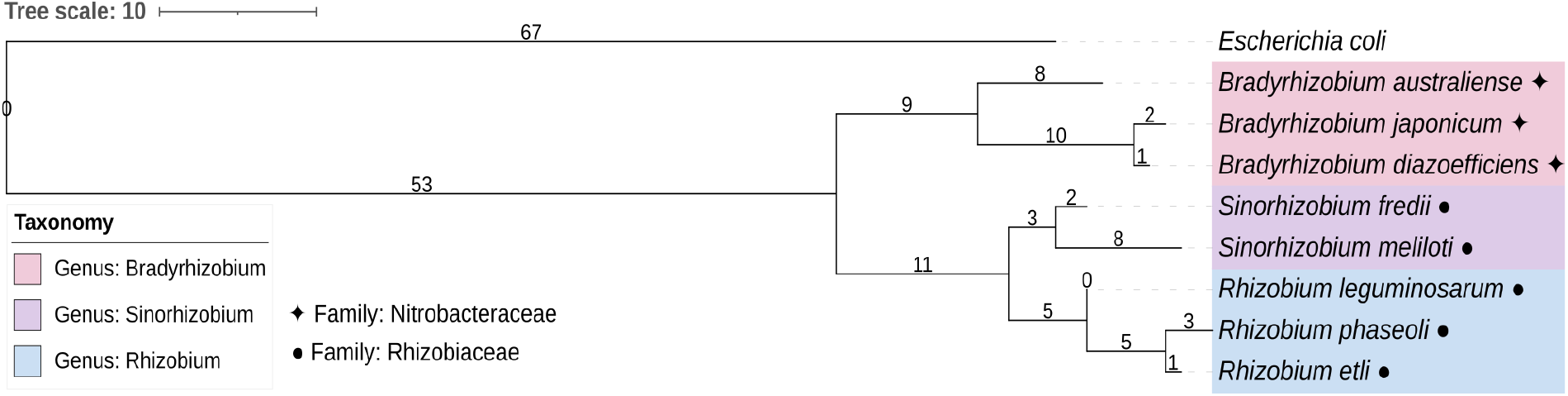
PuuR orthologous TFs in *rhizobia*. Phylogenetic tree inferred through Maximum Parsimony. Values on branches indicate phylogenetic distances. Color indicates genus, symbols indicate family of bacterial species.

## DISCUSSION

In this study, we identified TFs from *E. coli* whose function is conserved in *R. phaseoli*. To support our claim, we showed that known functional motifs are highly conserved, particularly DNA-binding motifs. By scanning the *R. phaseoli* genome, we identified TF-gene interactions and singled out PuuR-*nodI*, the one supported by gene expression, sequence homology, and functional evidence of being involved in symbiosis. The TRN presented here for *R. phaseoli*, although a valuable first of its kind among rhizobia, is an underestimation of the true network for various reasons: (1) it inherits the knowledge gaps present in *E. coli*, including the lack of annotated DNA-binding regions motifs and PSSMs for some TFs. Of the 75 oTFs, only 58 have an annotated DNA-binding region motif, and of the 49 oTFs whose DNA-binding motif passes the most stringent cut-off, only 25 have PSSMs. (2) the gene expression dataset used in this study to identify genes involved in the symbiotic process was obtained from a very specific point in time during the symbiotic process where *nod* genes were differentially expressed, *nif* genes to a lesser degree, and *fix* genes were not, indicative of the beginning of the symbiotic life cycle of *R. phaseoli* (5). To further illustrate this, the previously reported FixK⟶*fixN* interaction in *Rhizobium etli* (10) was recovered in this study at the sequence level, but *fixN* was not identified as differentially expressed. Finally, (3) our identification of TGs only considered the first gene of the operon. Future discovery of operon limits in *R. phaseoli* will provide a more comprehensive TRN. Further research to extend and add functional evidence for *R. phaseoli* TRN should take into account datasets obtained from different time points of the symbiosis and under a wide variety of conditions. The latest version of RegulonDB (33) provides publicly available ChIP-seq datasets that can be used to predict *de novo* TFBS for oTFs lacking a PSSM. The predicted TFBSs, although less reliable than PSSMs, can be used to scan upstream sequences in the *R. phaseoli* genome and therefore predict TGs for TFs lacking a PSSM.

PuuR in *E. coli* is known to bind putrescine, a polyamine (34, 35), which in cell plants has been implicated in protection against salt stress by stabilizing RNA, DNA, proteins, and phospholipids (35). Previous work in *Phaseolus vulgaris* determined that not only the number but also the variety of polyamines (including putrescine, spermidine, spermine, and homospermidine) found in nodules is higher compared to leaves and roots (35), suggesting a functional role in these structures. The regulation of PuuR over *nodI*, a nodulation gene, and *nifU2*, a nitrogen fixation gene, suggests a role in both nodulation and nitrogen fixation (**Figure 8**). Looking closer into PuuR binding-sites, it binds the *nodI* promoter at positions -411 to -390, and *nifU2* from -54 to -33. It has been established that repressors bind from -200 to +50 from the TSS while transcriptional activators could bind from -400 to -1 (36), placing PuuR as an activator of *nodI*, and repressor of *nifU2*, potentially regulating the transition from nodulation to nitrogen fixation.

**Figure 8.**
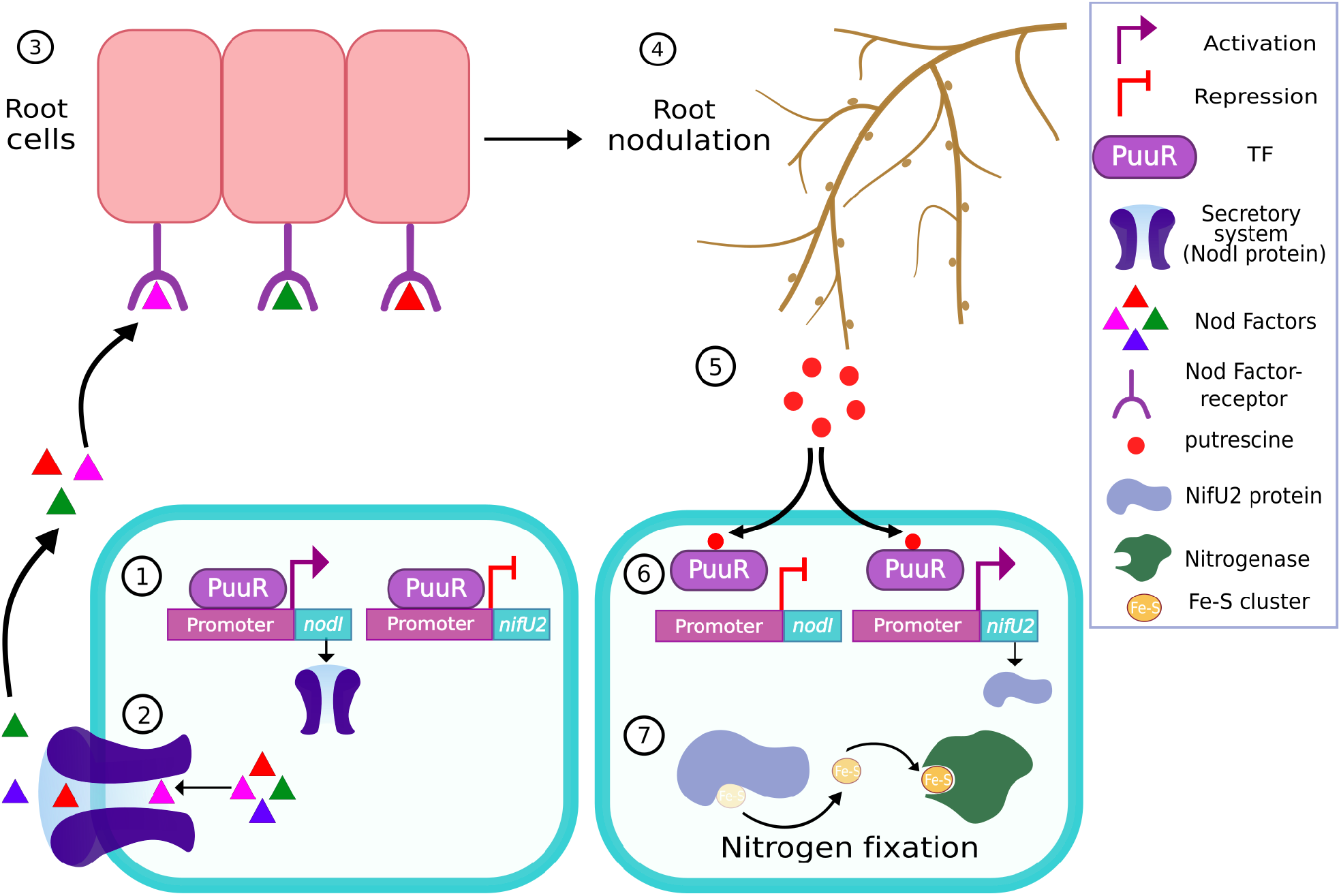
PuuR orthologous TF signaling model in *R. phaseoli*. PuuR binds DNA to activate *nodI* and suppress *nifU2. nodI*, a Nod Factors’ secretion system, facilitates Nod Factor export, triggering root cell signaling and root nodulation. Following nodule formation, putrescine released from plant roots is imported by *R. phaseoli*, binds intracellular PuuR, and the PuuR-putrescine complex unbinds from *nodI* and *nifU2* promoters. Halted repression of *nifU2* promotes its expression. *nifU2* encodes a supplier of precursors of [Fe–S] clusters, needed by the nitrogenase to fix nitrogen.

Taking together all the evidence presented here, we propose a model where PuuR activates *nodI* and represses *nifU2. nodI* is an NF secretion system (37, 38) whose expression promotes the export of NFs. NFs signal the root cells through NFs-receptors, triggering the nodulation process. Once the nodules are formed, putrescine is exuded from the plant roots and is imported by *R. phaseoli*. It binds PuuR causing it to unbind the *nodI* and *nifU2* promoter. *nodI is repressed and nifU2 is activated*, which codifies for a protein that supplies [Fe–S] clusters precursors for nitrogenase (39), and promotes nitrogen fixation.

Previous efforts to identify signaling molecules have been limited to flavonoids. Here, we propose a new signaling molecule along with its transcriptional response. Although we take advantage of the data available in *E. coli*, it is clear that the study of *R. phaseoli* and its symbiosis process requires the generation of high-resolution spatiotemporal data specific to this organism and its host plants. Deciphering the molecular cross-talk between the host plant and symbiont bacteria will allow us to improve, facilitate, and maintain the long-term colonization of crops, a crucial step for the development of sustainable agriculture strategies (6).

## MATERIALS AND METHODS

### Identification of orthologous TFs

Genomes were retrieved from NCBI (40): GCF_000005845.2 ASM584v2 for *Escherichia coli* str. K-12 substr. MG1655, GCF_000268285.2 RPHCH2410v2 for *Rhizobium phaseoli*, GCF_013114825.1 ASM1311482v1 for *Bradyrhizobium australiense*, GCF_004359355.1 ASM435935v1 for *Bradyrhizobium diazoefficiens*, GCF_013752735.1 ASM1375273v1 for *Bradyrhizobium japonicum*, GCF_002119845.1 ASM211984v1 for *Rhizobium etli*, GCF_004306555.1 ASM430655v1 for *Rhizobium leguminosarum*, GCF_003177055.1 ASM317705v1 for *Sinorhizobium fredii*, and GCF_000346065.1 ASM34606v1 for *Sinorhizobium meliloti*. Bidirectional Best Hits (41) were performed using BLASTp and the blastall software version 2.2.26. Only hits with e-value ≤ 1x10e-3, bit score ≥ 50, and identity percent ≥ 30 were considered. Cut-off values were selected based on previous literature (18).

#### Differentially expressed genes

Overexpressed genes from root exudates of maize, common bean, and milpa systems were considered from a Log Fold Change cut-off of 1, compared to an expression control in the absence of root exudates (26).

### Motif Analysis

For all the 75 oTFs, we retrieved the available annotated motifs from Ecocyc and observed whether they were conserved in the BBH alignment. First, BLASTp alignments were extracted by using blastall -m 0 option. Then, by using the motifs’ coordinates we pulled the motif sequence from the BLASTp alignment and computed the coverage and identity according to the BLAST glossary definition (42). Finally, we considered a motif conserved if the identity was ≥ 30% in both directions.

### Three-dimensional alignment

Protein structures were downloaded from the Alpha fold database (30) with the Uniprot identifiers “P0A9U6” for *E. coli* and “A0A0M3GCK6” for *R. phaseoli*. Subsequently, to align the DNA-binding region motifs of both TFs; coordinates 23-42 for *E. coli* and 41-60 for *R. phaseoli* as well as the option “jFATCAT (rigid)” were specified in the pairwise alignment tool.

### TRN assembly

PSSMs were obtained from RegulonDB (43). Upstream regions of *R. phaseoli* genes were defined from position -400 to +50, relative to the TSS, and were retrieved with the retrieve-sequence tool (44) of the RSAT suite using the parameters: -feattype CDS - type upstream -from -400 -to +50 -noorf -all. Upstream sequences were scanned with the PSSMs using the RSAT suite (44) tool matrix-scan (21) using the parameters: - consensus_name -pseudo 1 -decimals 1 -2str -origin end -bgfile 2nt_upstream-noorf_Rhizobium_phaseoli_Ch24-10_GCF_000268285.2_RPHCH2410v2-ovlp-1str.freq.gz - bg_pseudo 0.01 -return limits -return sites -return pval -return rank -lth score 1 -uth pval 1e-5.

### Gene annotation for nitrogen fixation and nodulation processes

*Rhizobium phaseoli* Ch24-10 genome annotation was downloaded with NZ_AHJU00000000 accession number and NZ_AHJU00000000.2 version from the NCBI Nucleotide database (40). Then, we searched for following regular expressions in the genome annotation file: “/gene=\”nod[A-Z]*[0-9]*[a-z]*” for *nod* genes, “/gene=\”nif[A-Z]*[0-9]*[a-z]*” for *nif* genes, and “/gene=\”fix[A-Z]*[0-9]*[a-z]*” for *fix* genes. Finally, the locus tag and protein ID were retrieved for each gene.

### Phylogenetic tree construction

The phylogenetic tree was inferred under the Maximum Parsimony optimization criterion. First, a multiple alignment of the sequences was performed using clustalω (45) with the following parameters: -pwgapopen=14 -pwgapext=0.2 - gapopen=14 -gapext=0.2. Then, we used the PAUP program version 4.0a (46) and provided as input the multiple alignment from clustalω. Additionally, the maximum parsimony optimization criterion, the bootstrap support values based on 1000 replicates, and the root of the tree were specified to PAUP. The outgroup method was used to root the three, and the *E. coli* protein was determined as the outgroup. Finally, the phylogenetic tree was visualized in the Interactive Tree Of Life (iTOL) online tool (https://itol.embl.de) (47).

## Data availability

All custom code and generated datasets can be found at: github.com/LedezmaTejeida-Lab/PutrescineAsSignalingMetabolite

## ACKNOWLEDGEMENTS

The authors thank Alfredo Hernández for server technical support, Alejandra Medina-Rivera for prompt RSAT support, and Enrique Merino-Perez and Luis Eduardo Servín-Garcidueñas for insightful discussions.

