## Supplementary material for "Putrescine acts as a signaling metabolite in the transition from nodulation to nitrogen fixation in *Rhizobium phaseoli*": Supplemetary Figures and Tables

[1] Centro de Ciencias Genómicas, Universidad Nacional Autónoma de México, Av. Universidad s/n, Cuernavaca 62210, Morelos, Mexico.

[2] Laboratorio Nacional de Ciencias de la Sostenibilidad (LANCIS), Instituto de Ecología, UNAM. Circuito Exterior s/n, junto al Jardín Botánico, Coyacán, Mexico City, 04510. Mexico.

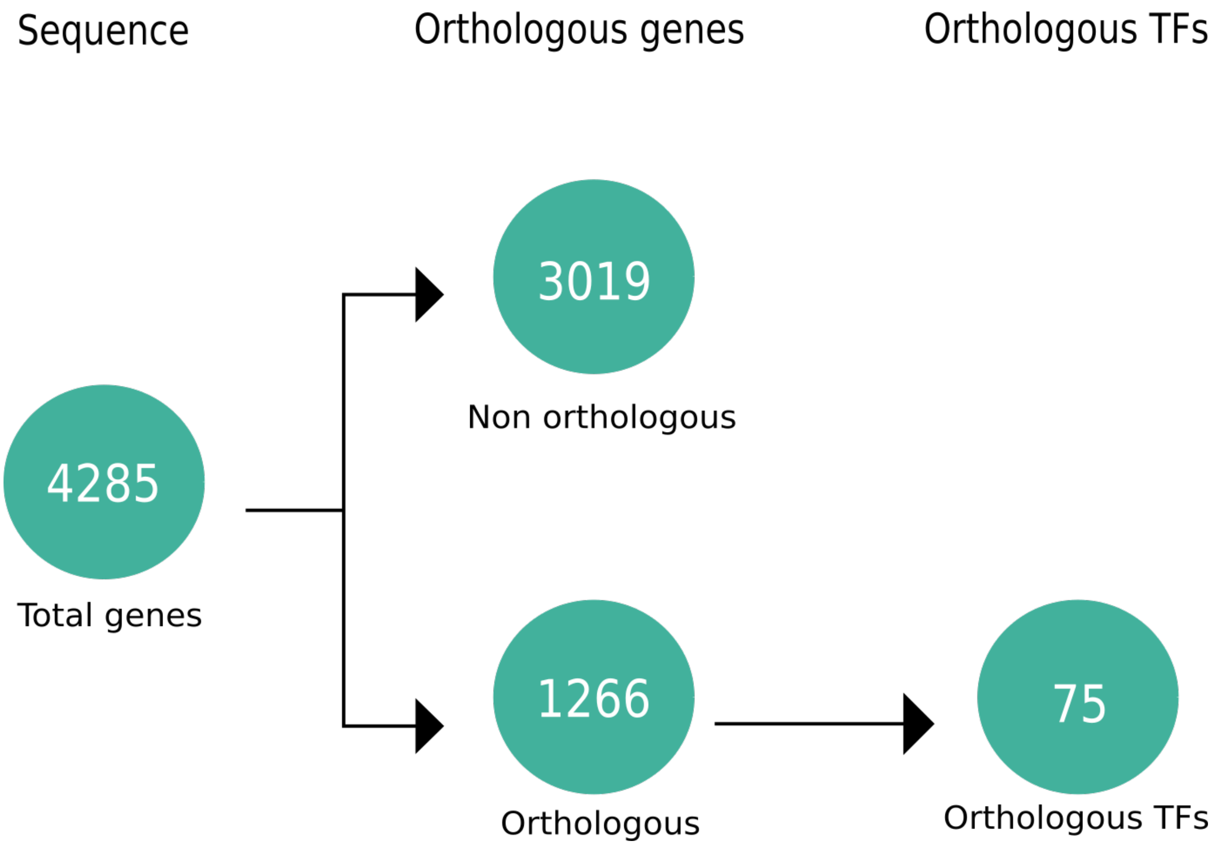


**Supplementary Figure 1. Orthologous genes searching**

The pipeline summarizes the number of genes filtered in each step. “Sequence” refers to the total number of *E. coli* coding genes. “Orthologous genes” refers to the number of orthologous and non-orthologous genes between *E. coli* and *R. phaseoli*. Additionally, “Orthologous TFs” refers to the number of orthologous TFs.

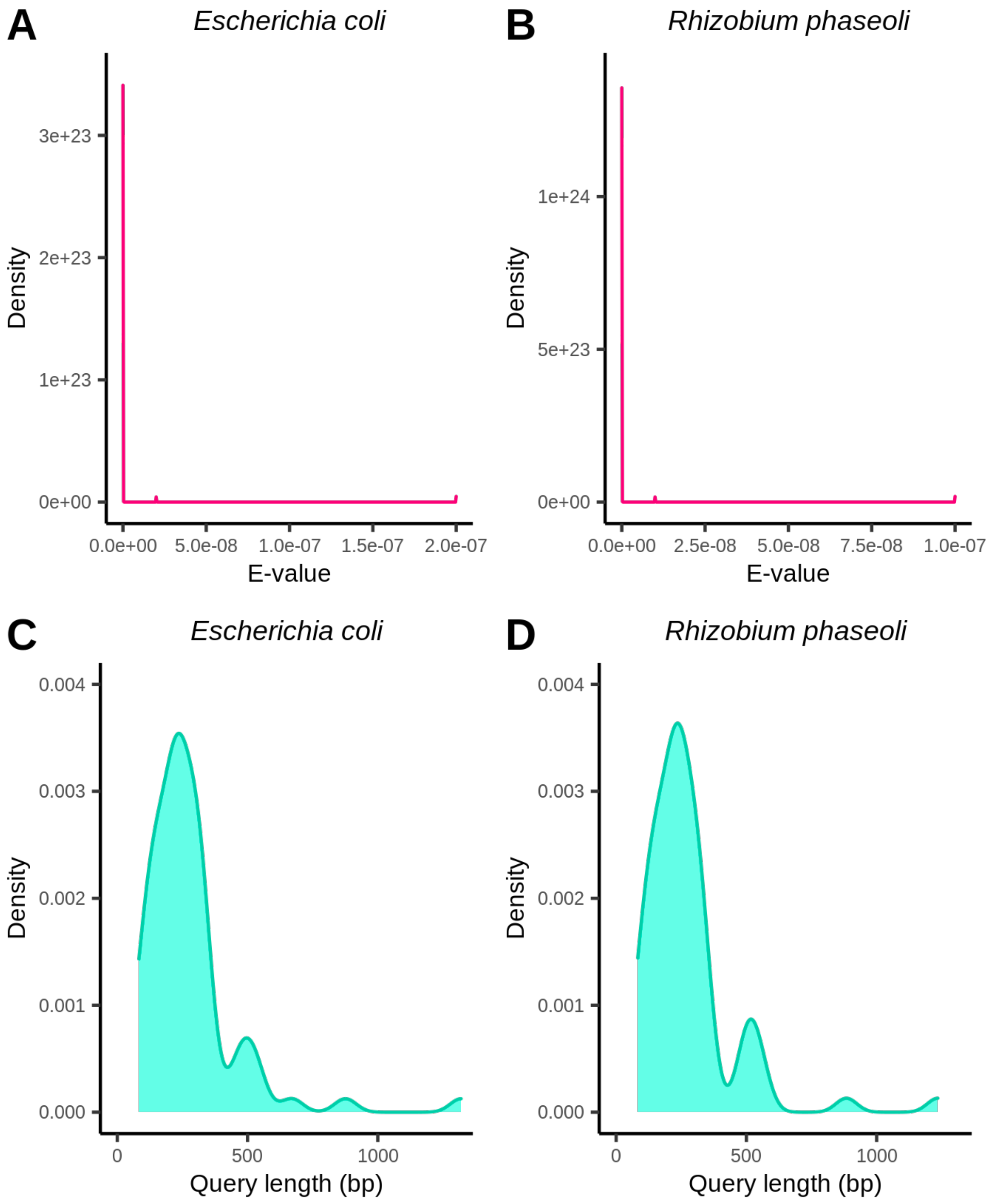


**Supplementary Figure 2. E-value and query length distributions**

E-value distributions of orthologous TFs, with (A) *E. coli* or (B) *R. phaseoli* genes as queries, are shown. C) and D) show the query length distribution of orthologous TFs with *E. coli* or *R. phaseoli* genes as queries, respectively.

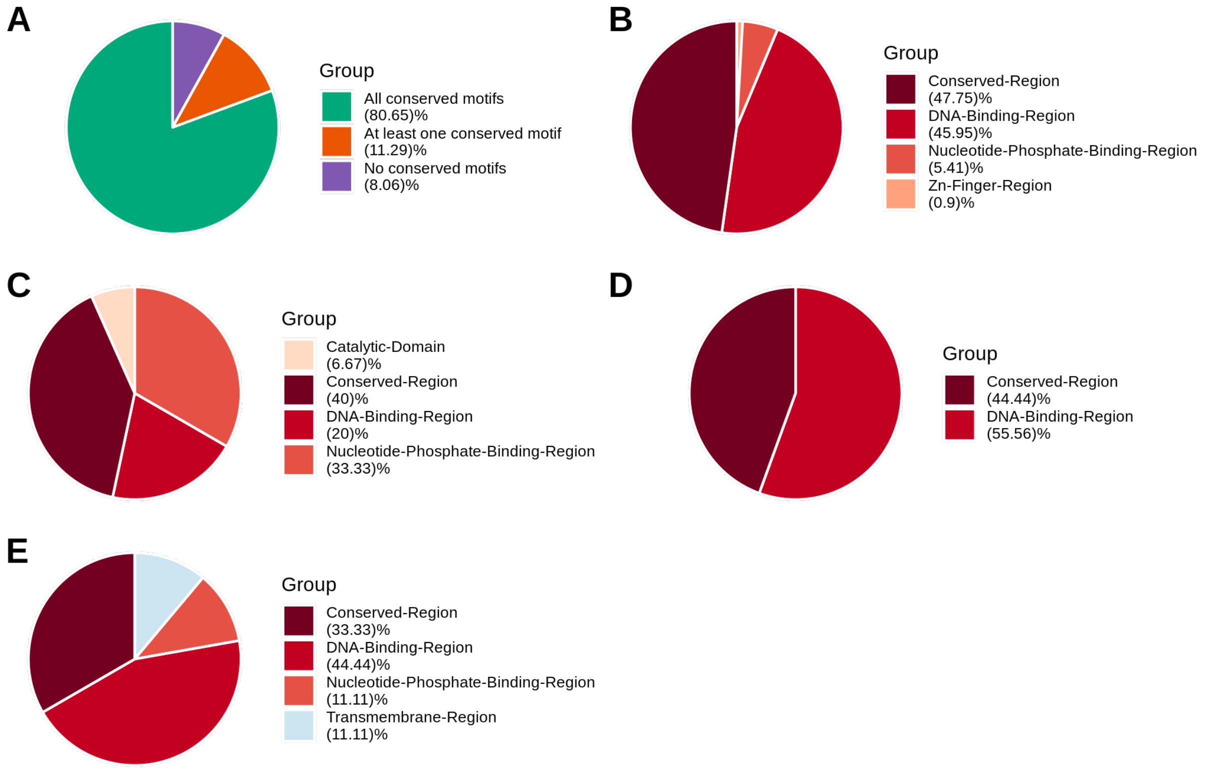


**Supplementary Figure 3. Motif conservation analysis**

A) Shows the proportion of oTFs having ‘All conserved motifs’, ‘At least one conserved motif’, and ‘No conserved motifs’. In addition, motif classes conserved in the ‘All conserved motifs’ B) and ‘At least one conserved motif’ C) categories are shown, as well as the classes of the no conserved motifs in the ‘At least one conserved motif’ D) or ‘No conserved motifs’ E) categories.

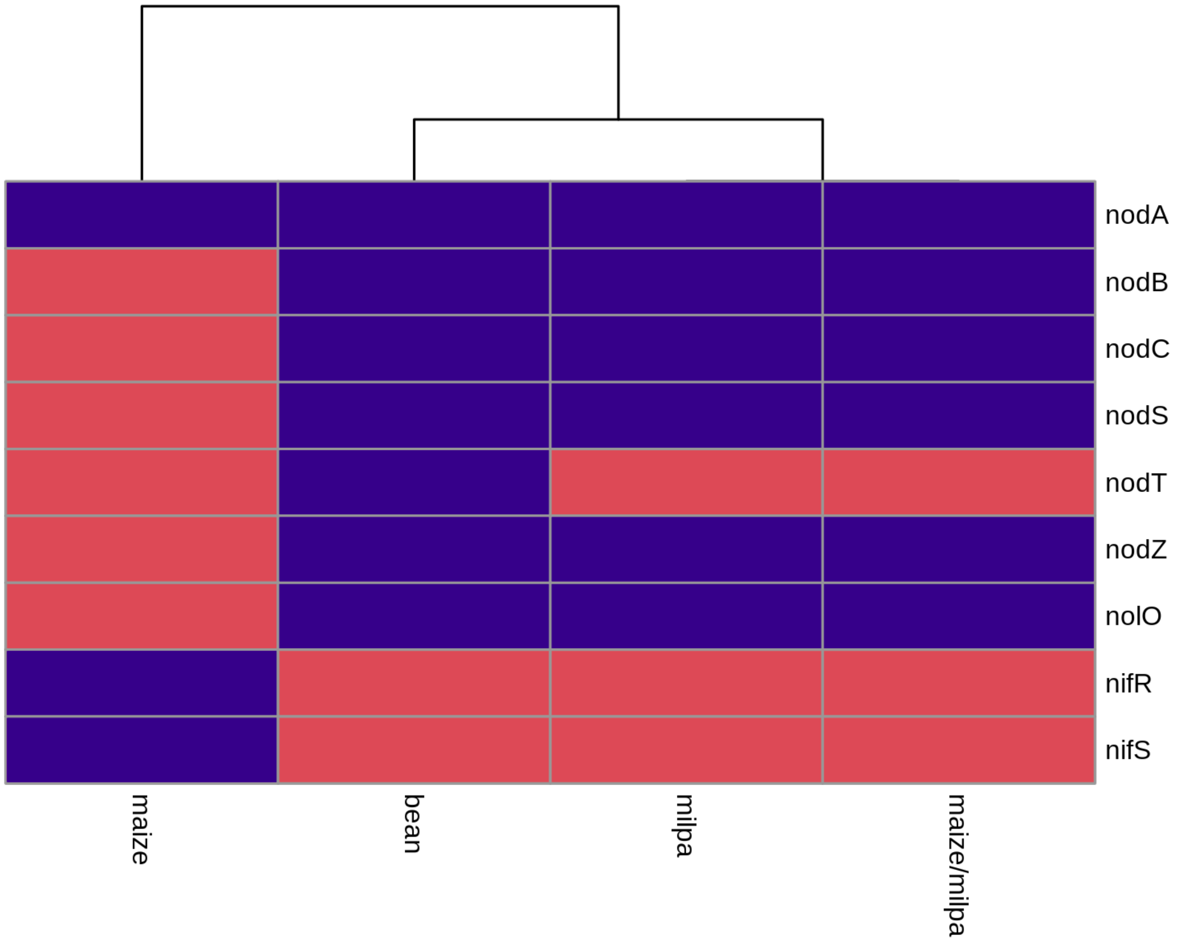


**Supplementary Figure 4. Evidence of function and gene expression**

Heatmap showing the presence (blue) or absence (red) of overexpressed symbiosis genes in the RNA-seq dataset. On the x-axis, the conditions under which the analysis was performed are indicated. On the y-axis, the names of the nine nodulation-related genes are shown.

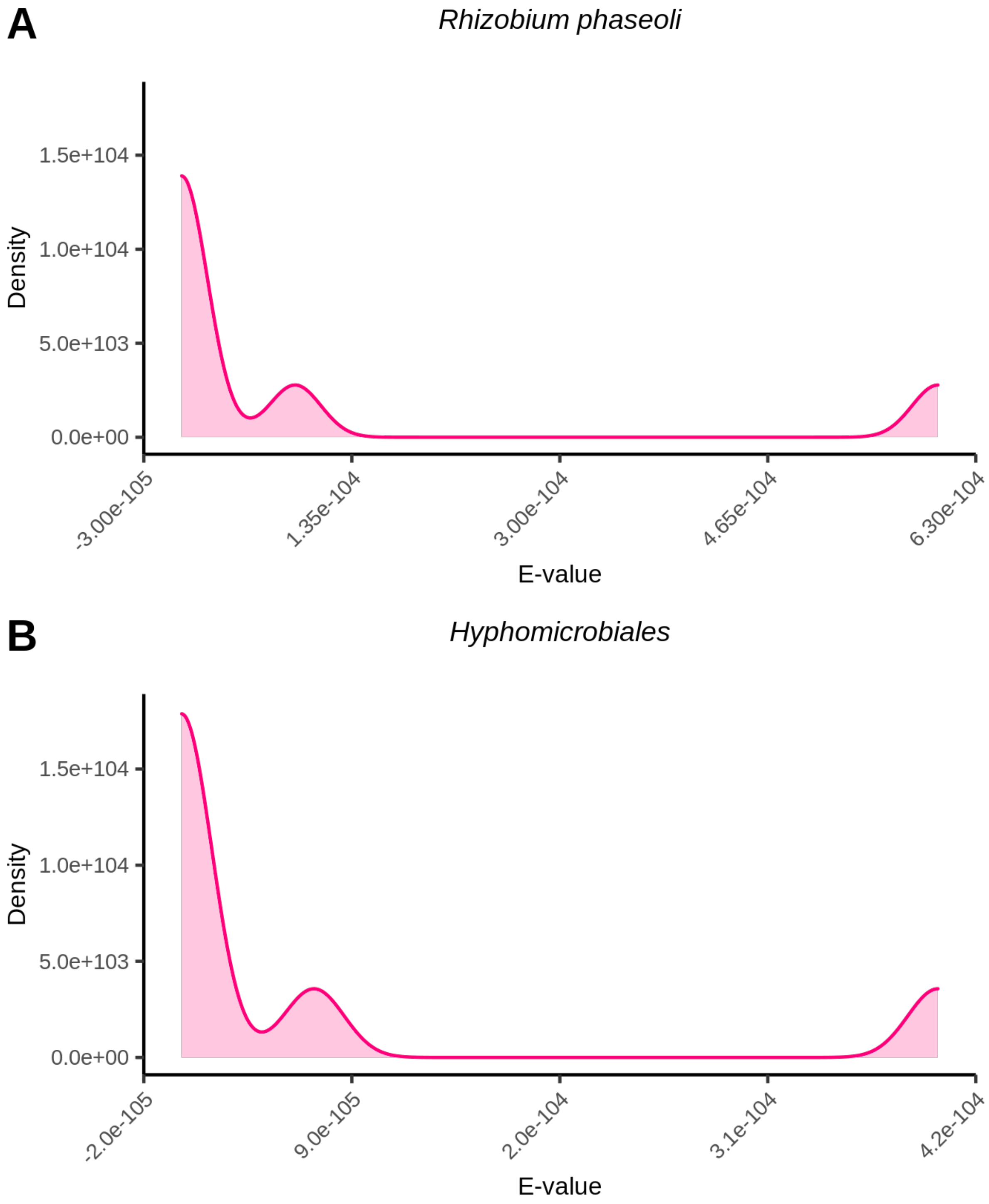


**Supplementary Figure 5. E-value distribution of PuuR oTFs**

E-value distributions of PuuR oTFs, with (A) *R. phaseoli* or (B) *Hyphomicrobiales* genes as queries, are shown.

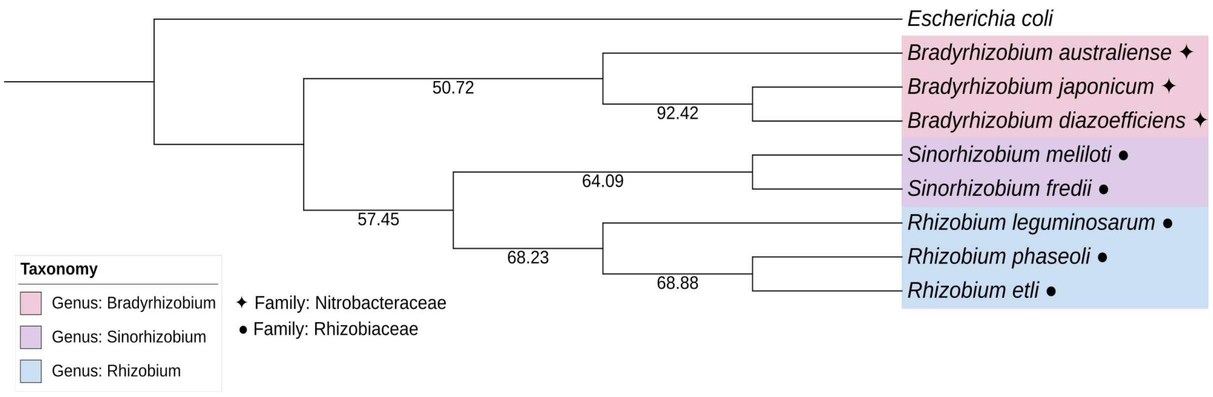


**Supplementary Figure 6. Bootstrap support of the phylogenetic tree**

Phylogenetic tree inferred under the criterion of Maximum Parsimony. The numbers on the branches indicate the bootstrap support for each partition after 1000 replicates. We highlighted with the same color the bacteria belonging to the same genus and indicated the family of bacteria (*Nitrobacteraceae* and *Rhizobiaceae*) with symbols.

| Motif | Description |
| --- | --- |
| Alpha-Helix-Region | Regions of a protein that form alpha-helical secondary structure motifs. |
| Beta-Strand-Region | Regions of a protein that form beta-strand secondary structure motifs. |
| Ca-Binding-Region | Calcium-binding region on a polypeptide. |
| Catalytic-Domain | Defines a domain that is associated with a particular catalytic activity. In a multifunctional protein, this feature can be used to associate different regions of a protein with different activities via the CATALYTIC-ACTIVITY slot. |
| Coiled-Coil-Region | A protein segment denotes the regions of a coiled-coil super-secondary structural motif. |
| Conserved-Region | Describes protein regions that share a high degree of sequence homology with similar regions in other proteins, but which cannot be definitively classified into any of the other feature classes. This class can include both homology motifs and larger conserved regions. |
| DNA-Binding-Region | DNA-binding region on a polypeptide. |
| Intramembrane-Region | A region of a polypeptide that is embedded in a membrane, but does not cross the membrane boundary on both sides. |
| Nucleotide-Phosphate-Binding-Region | Nucleotide-phosphate-binding region on a polypeptide |
| Protein-Structure-Region | Segments of a protein that are involved in protein structure levels above the primary structure, such as the beta strands or coiled coils. |
| Transmembrane-Region | A feature of most intrinsic proteins of plasma or vesicular membranes; a polypeptide sequence of about seven residues if beta-sheet, up to 22 residues if alpha-helix, that connects extracellular to intracellular domains, joined by the extended polypeptides on the cytoplasmic and external or vesicular sides. Receptor and ion channels have between one and twelve such domains. |
| Zn-Finger-Region | A motif of which seven or more may appear in a DNA-binding protein. Each is characterized by two closely spaced cysteine and two histidine residues that serve as ligands for a single $Zn^{2+}$. When bound to $Zn^{2+}$ they form a module from which protrudes amino acid side chains that interact with the bases partially exposed to the DNA major groove. Since in the area of contact, there is a limited and local parallel of DNA and protein sequences, the occurrence of the contacting amino acid residues in the protein sequence falls into a regularity, or register, that creates in the protein a syllabicity. |

**Supplementary Table 1. TF’s motif description**

Description of the motifs retrieved from Ecocyc (1).

| Gene name | Locus-tag | Protein ID |
|---|---|---|
| *fixA* | RPHASCH2410_PC00145 | KKZ84384.1 |
| *fixB* | RPHASCH2410_PC00150 | KKZ84385.1 |
| *fixC* | RPHASCH2410_PC00155 | KKZ84386.1 |
| *fixG* | RPHASCH2410_PC00515 | KKZ84444.1 |
| *fixGa* | RPHASCH2410_PA00890 | KKZ85303.1 |
| *fixH* | RPHASCH2410_PC00510 | KKZ84443.1 |
| *fixHa* | RPHASCH2410_PA00885 | KKZ85302.1 |
| *fixI* | RPHASCH2410_PC00505 | KKZ84442.1 |
| *fixIa* | RPHASCH2410_PA00880 | KKZ85301.1 |
| *fixJ* | RPHASCH2410_CH07435 | KKZ88987.1 |
| *fixKa* | RPHASCH2410_PA00915 | KKZ85308.1 |
| *fixLa* | RPHASCH2410_PA00920 | KKZ85309.1 |
| *fixLch* | RPHASCH2410_CH07430 | KKZ88986.1 |
| *fixN* | RPHASCH2410_PC00535 | KKZ84448.1 |
| *fixNa* | RPHASCH2410_PA00910 | KKZ85307.1 |
| *fixO* | RPHASCH2410_PC00530 | KKZ84447.1 |
| *fixOa* | RPHASCH2410_PA00905 | KKZ85306.1 |
| *fixP* | RPHASCH2410_PC00520 | KKZ84445.1 |
| *fixPa* | RPHASCH2410_PA00895 | KKZ85304.1 |
| *fixQ* | RPHASCH2410_PC00525 | KKZ84446.1 |
| *fixQa* | RPHASCH2410_PA00900 | KKZ85305.1 |
| *fixS* | RPHASCH2410_PC00500 | KKZ84441.1 |
| *fixSa* | RPHASCH2410_PA00875 | KKZ85300.1 |
| *fixX* | RPHASCH2410_PC00160 | KKZ84387.1 |
| *fnrN* | RPHASCH2410_PC00105 | KKZ84376.1 |
| *fnrNch* | RPHASCH2410_CH20730 | KKZ85764.1 |
| *nfeD* | RPHASCH2410_PA00455 | KKZ85217.1 |
| *nifA* | RPHASCH2410_PC00165 | KKZ84388.1 |
| *nifB* | RPHASCH2410_PC00170 | KKZ84389.1 |
| *nifD1* | RPHASCH2410_PC00080 | KKZ84371.1 |
| *nifD2* | RPHASCH2410_PC00605 | KKZ84461.1 |
| *nifD3* | RPHASCH2410_PC00360 | KKZ84420.1 |

| | | |
|---|---|---|
| *nifE* | RPHASCH2410_PC00070 | KKZ84369.1 |
| *nifH1* | RPHASCH2410_PC00085 | KKZ84372.1 |
| *nifH2* | RPHASCH2410_PC00610 | KKZ84462.1 |
| *nifH3* | RPHASCH2410_PC00355 | KKZ84419.1 |
| *nifK1* | RPHASCH2410_PC00075 | KKZ84370.1 |
| *nifK2* | RPHASCH2410_PC00600 | KKZ84460.1 |
| *nifN* | RPHASCH2410_PC00065 | KKZ84368.1 |
| *nifQ* | RPHASCH2410_PC00350 | KKZ84418.1 |
| *nifR* | RPHASCH2410_CH02630 | KKZ89515.1 |
| *nifS* | RPHASCH2410_PC00135 | KKZ84382.1 |
| *nifSch* | RPHASCH2410_CH21455 | KKZ85545.1 |
| *nifU* | RPHASCH2410_PC00130 | KKZ84381.1 |
| *nifU1* | RPHASCH2410_CH08880 | KKZ88239.1 |
| *nifU2* | RPHASCH2410_CH21170 | KKZ85489.1 |
| *nifW* | RPHASCH2410_PC00140 | KKZ84383.1 |
| *nifX* | RPHASCH2410_PC00060 | KKZ84367.1 |
| *nifZ* | RPHASCH2410_PC00180 | KKZ84391.1 |
| *nodA* | RPHASCH2410_PC00625 | KKZ84464.1 |
| *nodB* | RPHASCH2410_PC00470 | KKZ84437.1 |
| *nodC* | RPHASCH2410_PC00465 | KKZ84436.1 |
| *nodD1* | RPHASCH2410_PC00435 | KKZ84431.1 |
| *nodD2* | RPHASCH2410_PC00685 | KKZ84474.1 |
| *nodD3* | RPHASCH2410_PC00695 | KKZ84476.1 |
| *nodI* | RPHASCH2410_PC00450 | KKZ84434.1 |
| *nodJ* | RPHASCH2410_PC00445 | KKZ84433.1 |
| *nodL* | RPHASCH2410_CH08185 | KKZ88100.1 |
| *nodN* | RPHASCH2410_CH03920 | KKZ88303.1 |
| *nodS* | RPHASCH2410_PC00460 | KKZ84435.1 |
| *nodT* | RPHASCH2410_CH16550 | KKZ86733.1 |
| *nodW* | RPHASCH2410_PD03930 | KKZ84060.1 |
| *nodX* | RPHASCH2410_CH21465 | KKZ85547.1 |
| *nodZ* | RPHASCH2410_PC00485 | KKZ84438.1 |
| *nolL* | RPHASCH2410_PC00780 | KKZ84490.1 |

| | | |
|---|---|---|
| *nolO* | RPHASCH2410_PC00395 | KKZ84427.1 |

**Supplementary Table 2. Evidence of function.**

Gene name: displays the common name of the gene; Locus-tag: displays the unique identifier of the gene; Protein ID: displays the protein identifier according to the GenBank database.

| TF name | TG name |
|---|---|
| NarP | *nodD3* |
| ArgP, CytR, GcvA, UlaR | *nifA* |
| ExuR | *fixG* |
| FNR | *fixG(a), fixKa, fixN(a), fnrN(ch)* |
| GntR | *fixA* |
| PuuR | *nifU2* |

**Supplementary Table 3. Evidence of function and transcriptional regulation.**

The eleven genes with evidence of function and evidence of transcriptional regulation are shown. TF name: displays the TF name in *E. coli* that regulates the genes in the *R. phaseoli* TRN; TG name: displays the genes' names involved in nodulation or nitrogen fixation processes.

| Species Name | Locus-tag | Old Locus-tag | Protein ID (RefSeq) | Protein ID (GenBank) | Refseq Annotation | GenBank Annotation |
|---|---|---|---|---|---|---|
| *Rhizobium etli* | NXC12_RS31300 | NXC12_PE00602 | WP_020919480.1 | ARQ14196.1 | product="cupin domain-containing protein" | gene="puuR"; product="XRE family transcriptional regulator protein PuuR" |
| *Rhizobium leguminosarum* | ELH04_RS26215 | ELH04_26215 | WP_018245416.1 | TBE42178.1 | product="cupin domain-containing protein" | product="cupin domain-containing protein" |
| *Sinorhizobium meliloti* | SM2011_RS31170 | SM2011_c04387 | WP_010970549.1 | AGG75891.1 | product="cupin domain-containing protein"; function="small molecule metabolism" | product="Putative aldehyde dehydrogenase" |
| *Sinorhizobium fredii* | AOX55_RS18050 | AOX55_00003726 | WP_014330317.1 | AWM26956.1 | product="cupin domain-containing protein"; function="Predicted transcriptional regulators" | product="Putrescine utilization regulator" |
| *Bradyrhizobium diazoefficiens* | Bdiaspc4_RS11335 | Bdiaspc4_11360 | WP_011085008.1 | QBP21049.1 | product="cupin domain-containing protein" | product="cupin domain-containing protein" |
| *Bradyrhizobium japonicum* | HXT67_RS09810 | NA | WP_014497670.1 | NA | product="cupin domain-containing protein" | NA |
| *Bradyrhizobium australiense* | HCN58_RS33310 | HCN58_33435 | WP_028347208.1 | NOJ44393.1 | product="cupin domain-containing protein" | prpduct="cupin domain-containing protein" |

**Supplementary Table 4. *Hyphomicrobiales* order bacteria**

Each row corresponds to a bacteria belonging to the *Hyphomicrobiales* order. Species name: Genus and Species name of the bacteria; Locus-tag: Unique gene identifier of the gene*;* Old Locus-tag: Locus-tag in previous annotations; ProteinID (RefSeq): Protein identifier according to RefSeq database; Protein ID (GenBank): Protein identifier according to GenBank database; Refseq Annotation: Gene annotation according to Refseq database; GenBank Annotation: Gene annotation according to GenBank database.
